# Activation of Adipocyte mTORC1 Increases Milk Lipids in a Mouse Model of Lactation

**DOI:** 10.1101/2021.07.01.450596

**Authors:** Noura El Habbal, Allison C. Meyer, Hannah Hafner, JeAnna R. Redd, Zach Carlson, Molly C. Mulcahy, Brigid Gregg, Dave Bridges

**Affiliations:** Department of Nutritional Sciences, University of Michigan School of Public Health, Ann Arbor, Michigan, U.S.A.; Department of Pediatrics, University of Michigan Medical School, Ann Arbor, Michigan, U.S.A.

**Keywords:** Mammary glands, Milk composition, Adipocytes, mTORC1, Polyunsaturated Fatty Acids

## Abstract

Human milk is the recommended nutrient source for newborns. The mammary gland comprises multiple cell types including epithelial cells and adipocytes. The contributions of mammary adipocytes to breast milk composition and the intersections between mammary nutrient sensing and milk lipids are not fully understood. A major nutrient sensor in most tissues is the mechanistic target of rapamycin 1 (mTORC1). To assess the role of excess nutrient sensing on mammary gland structure, function, milk composition, and offspring weights, we used an Adiponectin-Cre driven *Tsc1* knockout model of adipocyte mTORC1 hyperactivation. Our results show that the knockout dams have higher milk fat contributing to higher milk caloric density and heavier offspring weight during lactation. Additionally, milk of knockout dams displayed a lower percentage of saturated fatty acids, higher percentage of monounsaturated fatty acids, and a lower milk ω6: ω3 ratio driven by increases in docosahexaenoic acid (DHA). Mammary gland gene expression analyses identified changes in eicosanoid metabolism, adaptive immune function, and contractile gene expression. Together, these results suggest a novel role of adipocyte mTORC1 in mammary gland function and morphology, milk composition, and offspring growth.

## Introduction

Human milk is considered the optimal source of nutrition for infants, and exclusive breastfeeding is recommended during the first 6 months of life (1). Successful lactation requires the development and differentiation of the mammary glands in preparation for milk production and secretion (2, 3). The mammary gland is composed of several cell types including adipocytes, contractile muscles, and alveolar cells. Mammary adipocytes are necessary for proper gland development and structure (4, 5). The mammary adipocytes in close proximity to the alveolar epithelial cells are thought to provide primary lipids for milk production (6). Given their role in maturation, development, and function of the mammary gland, adipocytes are crucial for successful lactation.

Maternal obesity has increased from around 26% in 2016 to 29% in 2019 (7). The health of the offspring is highly influenced by intrauterine and early postnatal exposures (8). During early postnatal life which is a critical developmental window, maternal obesity can alter breastfeeding capacity and milk composition (9). Maternal obesity can delay the initiation of lactation (10) and reduce the average duration of breastfeeding (11). The probability of early termination of lactation at three months postpartum was 1.5 times higher for infants of obese mothers compared to lean mothers (12). A meta-analysis of nine studies showed that maternal obesity increased mature milk fat composition but did not alter milk lactose or protein (13). The fatty acid composition of milk collected from obese women also showed a higher ω6:ω3 ratio compared to milk collected from non-obese counterparts (14).

Obese subjects have increased activity of the mechanistic target of rapamycin complex 1 (mTORC1) in the visceral fat compartment (15). mTORC1 is a critical nutrient sensor and a major regulator of protein and lipid synthesis (16, 17). mTORC1 promotes lipogenesis and adipogenesis and inhibits lipolysis (16, 18). In the presence of anabolic signals like insulin, energy abundance, and amino acid availability, mTORC1 function is upregulated (19). Developmental hyperactivation of mTORC1 in the mammary epithelium impairs the development of non-lactating mammary glands (20). However, little is known about the role of adipocyte mTORC1 with respect to macronutrient synthesis and secretion during lactation (21).

To elucidate the effects of excess nutrient sensing on lactation, we used a genetic adipocyte *Tsc1* knockout model. We show that chronic mTORC1 activation in maternal adipocytes increases adipocyte number and volume in mammary glands, changes milk fat levels and milk fatty acid composition, reduces gene expression of adaptive immune cell markers in the mammary glands, and increases the weight of lactating offspring.

## Materials and Methods

### Animal Husbandry

All mice were purchased from The Jackson Laboratory. Mice were fed a normal chow diet (Lab Rodent Diet; 5L0D) with *ad libitum* access to food and water. To hyperactivate adipocyte mTORC1 and generate an adipose-specific *Tsc1* knockout, *Tsc1*^fl/fl^ mice (JAX stock #005680, RRID: IMSR_JAX:005680; (22)) were crossed with *Adipoq*-Cre mice expressing the adipocyte-specific constitutive Cre recombinase controlled by adiponectin gene promoter (JAX Stock #010803, RRID: IMSR_JAX:010803; (23)).

The parental strains (F0) for this experiment were 6-8-week-old male *Tsc1*^fl/fl^; Cre^Tg/+^ (referred to here as knockout) and *Tsc1*^fl/fl^; Cre^+/+^ (referred to here as wild-type) crossed with 6-8-week-old female *Tsc1*^fl/fl^; Cre^+/+^ and *Tsc1*^fl/fl^; Cre^Tg/+^, respectively. The offspring (F1) were thus also a combination of knockout and wild-type mice. The Cre driver was Adiponectin-Cre, which is expressed in all adipocyte lineages (brown, white, and mammary adipocytes; (24, 25)). There is currently no known Cre driver that is specific to mammary adipocytes. All animal procedures were carried out in accordance with the National Institute of Health Guide for the Care and Use of Laboratory Animals and was approved by the University of Michigan Institutional Animal Care and Use Committee prior to the work being performed.

After 16 days of mating, male breeders (F0) were removed from the cage to avoid the occurrence of a second pregnancy. We checked for litters on a daily basis after 2.5 weeks of mating. The number of offspring born (F1) was recorded to determine maternal fertility and offspring viability. After delivery (delivery day denoted as postnatal day 0.5, PND0.5), the dams continued to have *ad libitum* access to food and water. At PND2.5, offspring were sexed by anogenital distance assessment, and litters were culled to four animals (2 females and 2 males, when possible) to normalize milk supply.

### Body Composition

Mice were weighed by dynamic weighing to capture accurate measurements using a digital scale (Mettler Toledo, ML6001T). For body composition assessment, mice were placed in the magnetic resonance imaging (MRI) tube and restrained during the magnetic resonance imaging measurement (EchoMRI, EchoMRI 1100). Fat, lean, free water, and total water mass (g) were recorded for each animal. Dams (F0) in all groups were weighed and underwent body composition assessment via MRI three times a week during pregnancy and lactation. On the day of delivery, dams were weighed and their body composition was assessed via MRI. At the end of our experiment, on PND16.5, all dams were weighed and underwent MRI prior to milk collection then were immediately euthanized. The offspring (F1) were weighed at PND0.5, 7.5, and 14.5. On PND16.5, the offspring were weighed and underwent body composition assessment via MRI then were immediately euthanized.

### Determining Milk Output Volume

At peak lactation (PND10.5; (26), we determined milk output volume for wild-type and knockout dams. To determine milk volume, we used the weigh-suckle-weigh technique (27). Briefly, we weighed the dam separately and the offspring in aggregate. The dam and offspring were then separated for two hours to allow the milk reserves of the dam to build. During the two-hour separation, the offspring were placed in a new cage and were kept warm using a heating pad under the cage. In the meantime, the dam remained in its initial cage with *ad libitum* access to normal chow diet and water. After the two-hour separation period, the dam was weighed and the aggregate weight of the offspring was measured, each for a second time. The offspring were then returned to the home cage and were allowed to nurse for one hour undisturbed. At the end of the nursing period, the dam was weighed again, and the aggregate weight of the offspring was measured for a third time. Milk volume was approximated as the weight change of the offspring or the dams after the one hour nursing period.

### Euthanasia and Tissue Collection

All dams were sacrificed using anesthetic gas inhalation (5% isoflurane via drop jar method) at PND16.5 after milk collection. We extracted thoracic, inguinal, and abdominal mammary glands. Briefly, the peritoneum was pulled away from the skin, and the inguinal and abdominal mammary glands (referred to here as lower mammary glands) were excised completely and weighed. The tissue from the left and right lower mammary glands were collected in 2mL Eppendorf tubes, snap frozen in liquid nitrogen, and later stored at −80°C for molecular assays. Portions of the thoracic mammary glands (referred to here as upper mammary glands) were fixed in 10% formalin, dehydrated in 70% ethanol, and later embedded in paraffin for histology and adipocyte assessment. Offspring of dams were sacrificed without tissue extraction at PND16.5 after body weight and composition measurements.

### Milk Collection

On PND16.5, we collected milk samples (∼0.5mL) from the nursing dams. To do this, after two hours of separation from the offspring, we anesthetized the dam by intramuscular injection of Ketamine/Xylazine (0.1275g/kg body weight) into the hindlimb muscle. Once the dam was fully anesthetized, we then injected oxytocin intramuscularly into the forelimb (2U/dam) to induce milk let-down. The dam’s nipples were manually squeezed to promote milk secretion, and the milk was collected into a 1.5 mL tube via suction using a 50mL conical vacuum apparatus. After milking was complete, the dam was immediately euthanized using isoflurane and the mammary glands were extracted.

### Milk Composition Assessment

Milk samples collected from wild-type and knockout dams on PND16.5 were assessed for fat content by the creamatocrit method using a microhematocrit centrifuge (28). Briefly, whole mouse milk was diluted four-fold with 1X PBS-EDTA solution. Milk samples were transferred into plain microhematocrit glass capillary tubes. The tubes were sealed from one end using Critoseal. The tubes were later placed in a microhematocrit centrifuge (Iris Sample Processing, StatSpin CritSpin M961-22). Samples were centrifuged for 120 seconds per cycle for a total of 8 cycles and a total spin time of 16 minutes. The capillary formed layers of non-fat milk and white fat. The length of the white fat layer was measured using a 150 mm dial caliper (General Tools, 6” Dial Caliper). The total volume of milk (non-fat and fat milk layers) was also measured in mm. Percentage of fat was determined with respect to the total milk volume.

Lipidomic analyses were done by the Michigan Regional Comprehensive Metabolomics Core. Milk samples were frozen at −80°C until analysis to prevent lipid hydrolysis and peroxidation. Samples were quickly thawed once for lipidomic analysis without undergoing multiple freeze-thaw cycles. Long chain fatty acid concentrations were determined following sample extraction, semi-purification and derivatization followed by fatty acid measurement by gas chromatography (GC) using an Agilent GC model 6890N equipped with flame ionization detector. Results were reported on 33 lipid classes from C14:0 to C24:1.

### Transcriptomic Analyses

Using the lower right mammary gland tissues collected from the dams on PND16.5, we assessed whole-transcriptome RNA expression using five wild-type and six knockout samples. RNA samples were prepared from the mouse tissues using the PureLink RNA Mini Kit (Invitrogen by ThermoFisher Scientific, catalog #12183025). Briefly, tissues were cut on dry ice to ∼50mg then homogenized and treated to collect purified RNA. The RNA was quantified using a nanodrop, and purity was verified by an Agilent Bioanalyzer. All samples had an RNA integrity number (RIN) higher than 7. Library preparation and next generation sequencing was conducted by the Advanced Genomics Core at the University of Michigan. Paired-end poly-A mRNA libraries were generated and sequenced to an average depth of 57M (range 46M-69M) reads per sample on Illumina NovaSeq 6000 (S4). Reads were aligned to the mouse reference genome GRCm38.p6 using Salmon v 1.3.0 (RRID: SCR _017036; (29)) with the gc-bias and validateMappings flags. Mapping efficiency was 54.8% (sample range 53-56.6%). Transcript-level data was reduced to gene-level data via tximeta v1.8.4 (30) and txiimport v1.18.0 (31) prior to analysis by DESeq2 v1.30.1 (32). To determine differentially expressed genes (DEGs), we evaluated 14242 genes, excluding those with low or no read counts. Full gene expression results are reported in Supplementary Table 1. For gene set enrichment analyses, we used ClusterProfiler v3.16 (RRID: SCR_016884; (33)) after ranking genes by fold change and analyzing relative to Gene Ontologies. Similarities between enriched gene sets were calculated by Jaccard distances. Gene set enrichment results are presented in Supplementary Table 2. Data are available from GEO at accession number GSE175620.

### Mammary Gland Histology and Adipocyte Assessment

Upper right mammary glands collected from wild-type and knockout dams were embedded in paraffin and Hematoxylin and Eosin (H&E) stained at the Rogel Cancer Center’s Tissue and Molecular Pathology Core. Slides were blindly assessed for adipocyte size and number using one slide per mouse. Using an EVOS XL Imaging System inverted fluorescent microscope (Invitrogen by ThermoFisher Scientific, catalog # AME3300), eight representative pictures per slide were taken at 10x magnification and covered the entire tissue area. Mammary gland adipocytes were quantified using ImageJ with Adipocyte Tools Macros Plugin (34). In analyzing our images, the parameter filters for adipocytes using the processing options were set at minimum of 40 pixels, maximum of 1000 pixels, and 30 dilates. The parameters for segmentation options were set at minimum of 600 pixels and maximum of 1500 pixels. Potential adipocytes that were blurry, cut off, or below the threshold were excluded from the assessment to maintain accurate measurements. Once these two parameters were set on the image, manual additions and deletions were performed to ensure adipocytes were properly identified. After accounting for all adipocytes, they were further analyzed using the ImageJ software to obtain area measurements. The calculated adipocyte numbers were normalized to the total mammary gland area that was imaged.

### Statistical Analyses

Statistical significance was designated at p<0.05 for this study. All statistical analyses were performed using R v4.0.2 (35). Data are presented graphically as mean +/− standard error of the mean. For longitudinal measurements including body composition, food intake, and offspring weight gain, data were analyzed using mixed linear models using lme4 v1.1-25 (36). We tested for sex-modification of all offspring outcomes involving both sexes and report these when significant. For pairwise testing, normality was assessed using Shapiro-Wilk tests followed by homoscedasticity using Levene’s test. Pending these results, appropriate parametric or non-parametric tests were done. All raw data, statistical analyses, and reproducible R code can be found at http://bridgeslab.github.io/TissueSpecificTscKnockouts/.

## Results

To understand how activation of mTORC1 in adipocytes affects lactation, we evaluated pregnant mice that were either wild-type or adipocyte *Tsc1* knockout. Virgin dams were mated with a male having the opposite genotype. The experimental timeline and mouse models are shown in Figure 1A-B.

**Figure 1:**
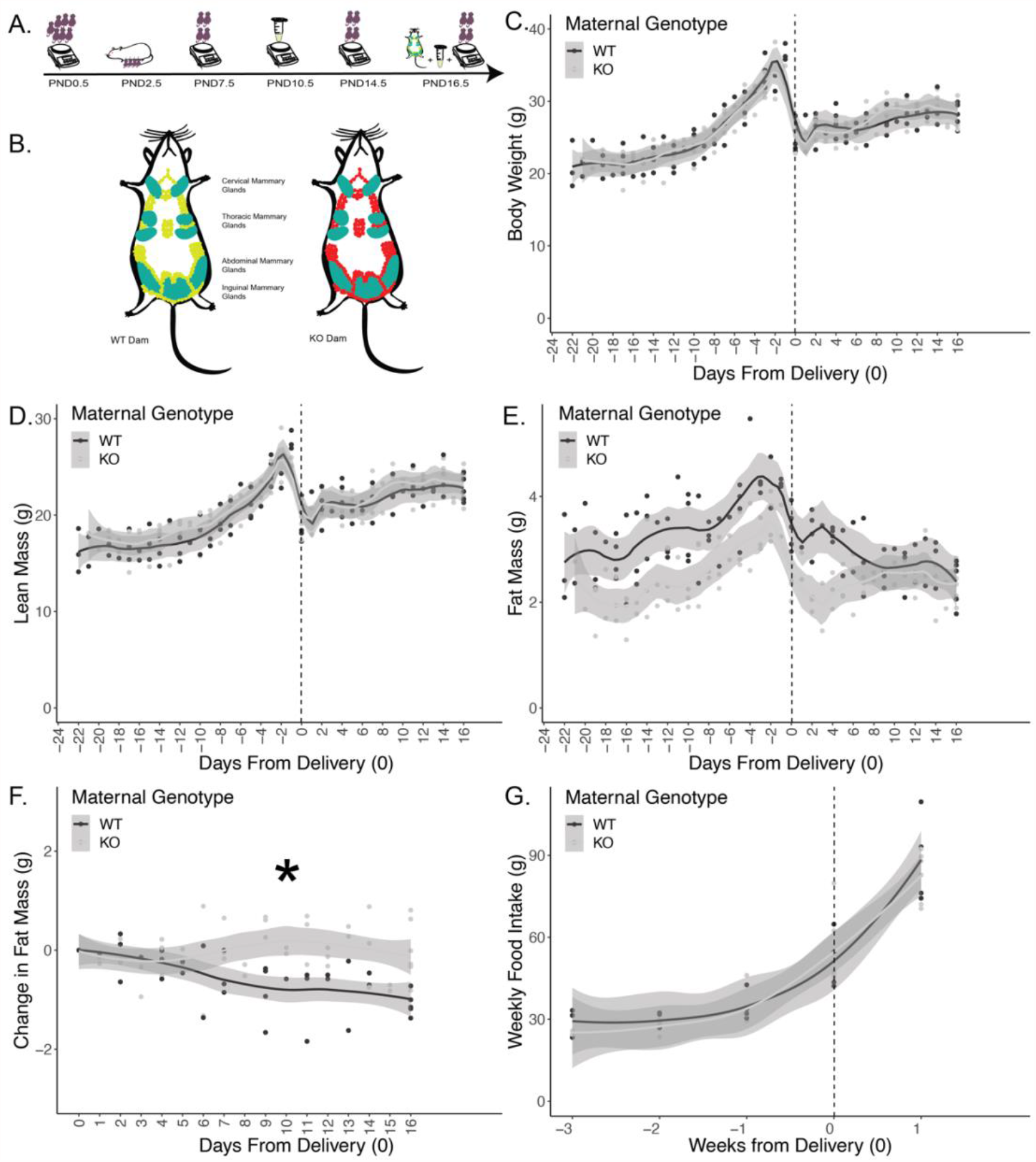
Experimental timeline and dam body composition. (A) Experimental timeline. Dams and offspring were monitored throughout lactation. Offspring were born and weighed at PND0.5. Litters were culled to 4 offspring (2 males and 2 females) on PND2.5. Milk volume measurements were conducted on PND10.5. Offspring were weighed on PND7.5 and PND14.5. PND16.5 marks the end of the experiment where the dam and offspring were weighed then euthanized, milk was collected from mammary glands of the dam, and mammary glands were excised for histological and molecular studies. Maternal body composition was measured on PND0.5 after delivery and three times a week thereafter until and including PND16.5. (B) Schematic showing mammary glands (in teal) and whole body adipocytes for wild-type (WT; in green) and knockout (KO; in red) dams with *Tsc1* deletion. (C) Maternal body weights. (D) Maternal lean mass. (E) Maternal fat mass. (F) Maternal change in fat mass postnatally from the day of delivery until PND16.5 reveals more fat mass gain during lactation in knockouts. (G) Weekly food intake. Asterisk indicates p<0.05 from mixed linear model. (n=11 dams; 5 wild-type and 6 knockout).

### Fat Mass Gain in Adipocyte *Tsc1* Knockout Mice during Lactation

Dam body weight and composition were assessed during pregnancy and lactation. Body weights were comparable between dams throughout the study (Figure 1C). Lean mass was also similar between adipocyte *Tsc1* knockout and wild-type dams (Figure 1D). Knockout dams had a slightly lower fat mass during pregnancy and during lactation (Figure 1E). While wild-type dams lost a small amount fat mass during lactation, knockout dams gained fat mass (Figure 1F, p<0.001). This was not explained by differences in food intake, as wild-type and knockout dams had similar food intake throughout pregnancy and lactation (Figure 1G).

### Adipocyte *Tsc1* Knockout Mice Have Smaller Mammary Glands with More and Larger Adipocytes

At sacrifice on PND16.5, we examined the inguinal and abdominal mammary glands. As shown in Figure 2A, knockout dams had a 21% reduction in mass of the right mammary glands (p=0.042) and a 29% reduction in mass of the left mammary glands (p=0.001) compared to the wild-type counterparts.

**Figure 2:**
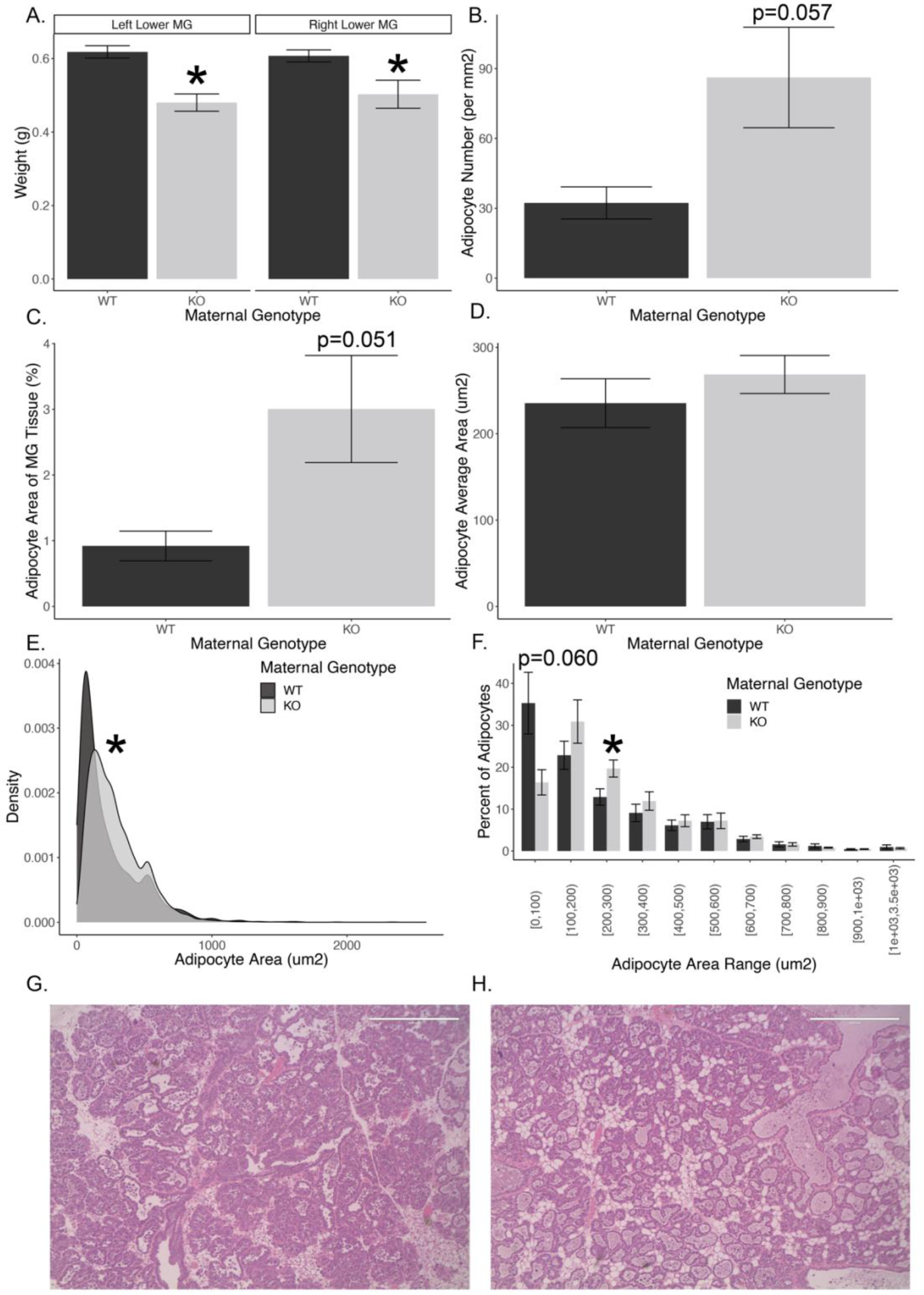
Mammary glands histology. (A) Inguinal and abdominal (lower) mammary gland (MG) weights. (B) Histological analysis showing increased number of adipocytes in adipocyte *Tsc1* knockout thoracic (upper) mammary glands. (C) Average area of the mammary gland comprised of adipocytes. (D) Mean adipocyte area. (E) Distribution of adipocyte area, cumulative over all images. (F) Percent of adipocytes by genotype and area range. For E, the asterisk indicates a significantly different distribution of adipocyte sizes by a Kolmogrov-Smirnoff test. (G) Representative images of a wild-type (G) or knockout (H) thoracic mammary gland sections. For B and C, mixed linear models were used with images as the random effect and genotype as the fixed effect. (n=11 dams, 8 images per dam, 174-3199 adipocytes per slide, 10533 total adipocytes for knockout dams and 3286 total adipocytes for wild-type dams).

Right thoracic mammary glands were fixed and stained for histological analyses to assess the number and area of adipocytes in the dams (Figure 2G-H). Adipocyte *Tsc1* knockout mammary glands had 63% more adipocytes compared to the wild-type dams (Figure 2B, p=0.057). We then assessed the adipocyte area as a fraction of the total mammary gland and found that in the knockout, adipocytes occupied three times the mammary gland area compared to the wild-type mammary adipocytes (Figure 2C, p=0.051).

The mean area for mammary adipocytes was not significantly different (Figure 2D, p=0.36), however, the distribution of adipocyte sizes was different. Adipocyte *Tsc1* knockout mammary adipocytes had a significantly wider variation in the distribution of adipocyte area (Figure 2E, p<0.001). Knockout mammary glands had 52% more larger sized adipocytes (200-300 µm^2^) compared to wild-type (Figure 2F, p=0.039). Similarly, *Tsc1* knockout mammary glands had 46% fewer adipocytes in the smallest range (0-100 µm^2^) compared to wild-type adipocytes (Figure 2F, p=0.060). Our results show histological differences where adipocyte *Tsc1* knockout dams have more adipocytes, more larger adipocytes, and a higher percentage of total mammary gland area comprised of adipocytes, despite having smaller mammary glands overall.

### Offspring Born to Adipocyte *Tsc1* Knockout Dams are Heavier During Peak Lactation

The average litter size across genotypes was similar (Figure 3A). Litters were culled to four offspring per dam at PND2.5 to normalize milk supply. There was no significant difference in offspring weight at birth (PND0.5; Supplementary Figure 1A). At PND7.5, after adjusting for sex, offspring born to knockout dams were 7% heavier than offspring born to wild-type dams (Figure 3B, p=0.010 from a 2×2 ANOVA). Female offspring born to knockout dams were 9% heavier than females born to wild-type dams (Figure 3B, p=0.044). Weights of male offspring born to knockout dams were 5% heavier than males born to wild-type dams although this did not reach statistical significance (Figure 3B, p=0.14). At PND14.5 and PND16.5, there were no weight differences between groups or sexes (Supplementary Figure 1B). Our results show that the offspring of knockout dams are heavier during the first week of life when they are solely reliant on lactation for nutrient acquisition. At later time points, we hypothesize that the weights converge between genotypes because the offspring are eating more chow-based food and rely less on maternal lactation.

**Figure 3:**
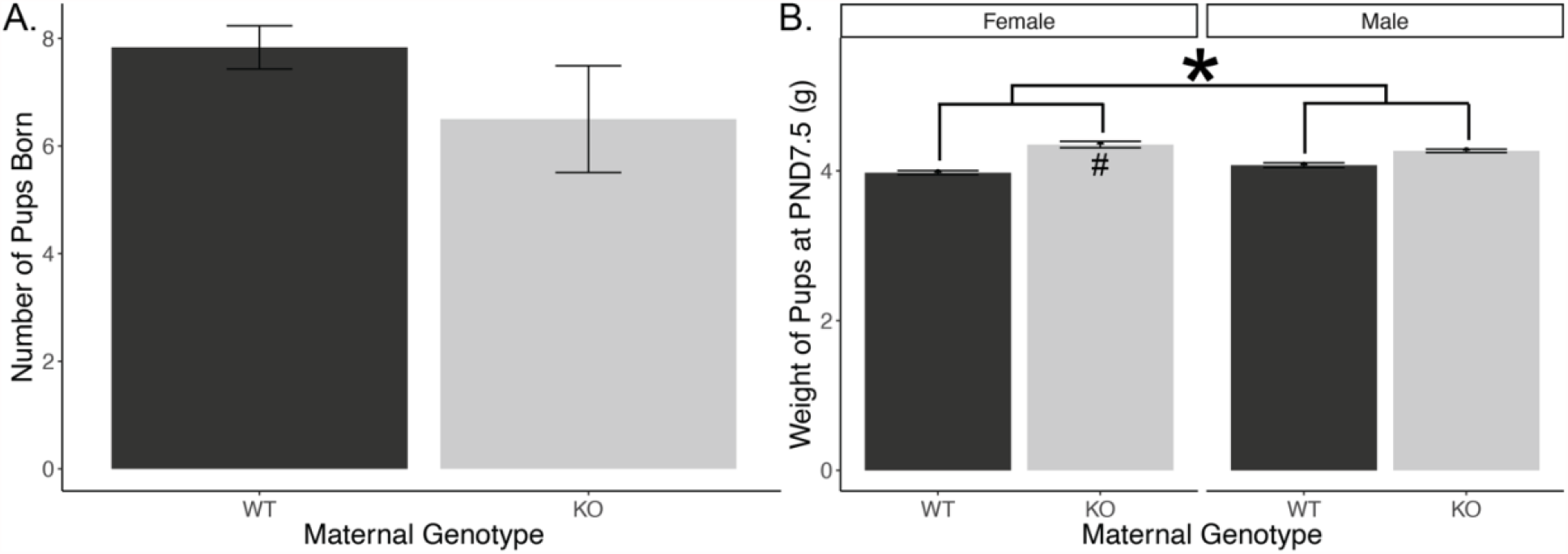
Litter size and offspring weight. (A) Average number of offspring born to wild-type and knockout dams per their single litters. (B) Weights of male and female offspring of wild-type and knockout dams at PND7.5. Asterisk indicates a significant effect of maternal genotype from 2×2 ANOVA; number sign indicates significance within female offspring from a pairwise t-test. (n=44 offspring from 11 dams).

### Adipocyte *Tsc1* Knockout Dams Produce Similar Volumes of Milk but with a Higher Milk Fat Percentage and Increased Desaturation of Milk Fatty Acids

Due to the differences in offspring weight and mammary gland size and histology, we calculated the mass of milk produced per dam at PND10.5. Milk output of the dams was not significantly different between groups (Supplementary Figure 2A-B). This suggests that the differences we saw in offspring weight were not driven by increased milk output in the knockout dams. This prompted us to further evaluate the milk fat composition. Milk was collected from dams at PND16.5. Total fat analysis using the creamatocrit technique revealed that milk of adipocyte *Tsc1* knockout dams had a 34% higher fat percentage than milk of wild-type dams (Figure 4A, p=0.024). This finding shows that the increased fat content in the milk of knockout dams could be the main driver in increasing offspring weight due to increased milk caloric density.

**Figure 4:**
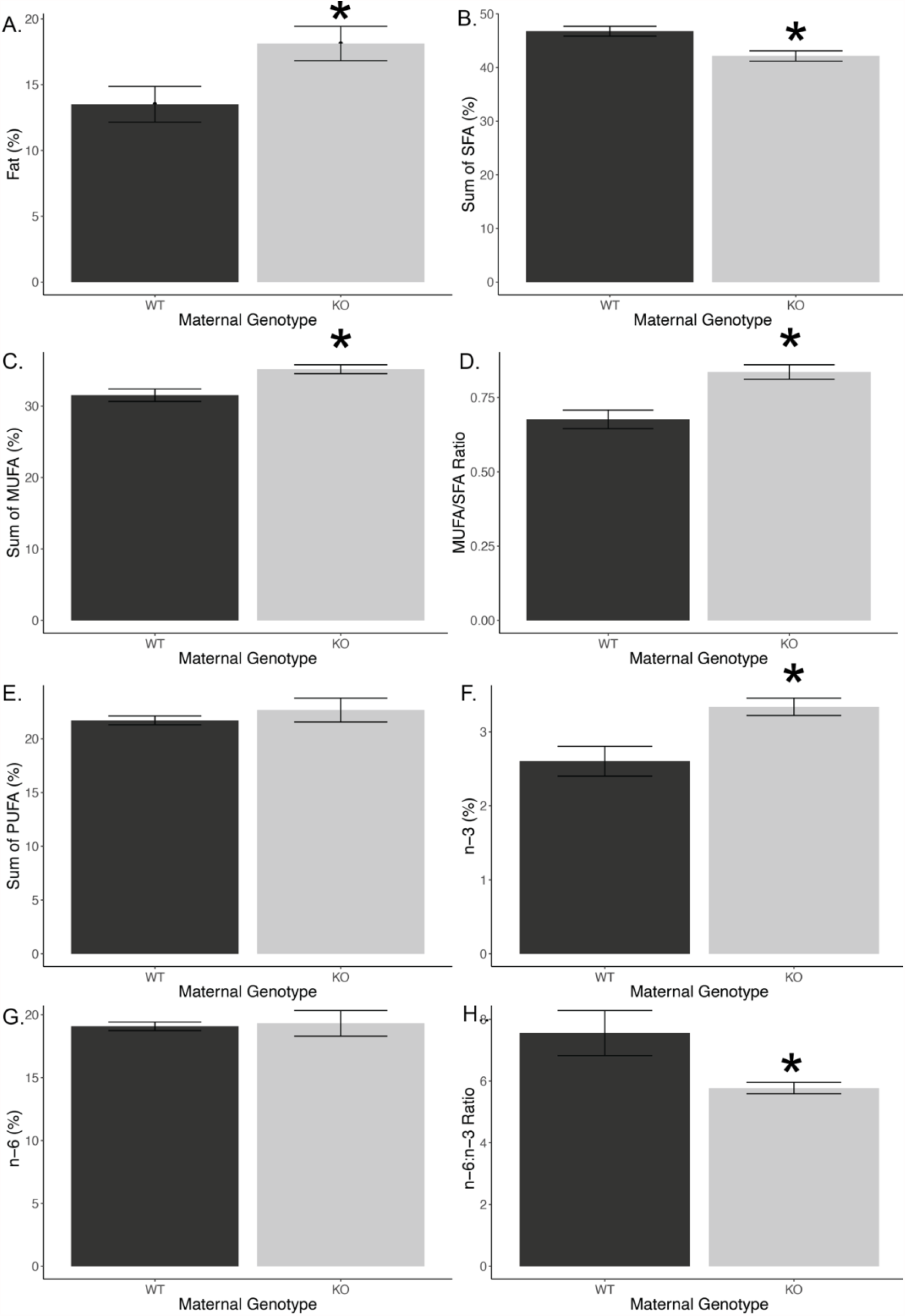
Milk fat analysis. (A) Average percent fat composition of milk from knockout and wild-type dams. (n=21 dams). (B) Percentage of saturated fatty acids (SFA) in milk. (C) Percentage of monounsaturated fatty acids (MUFA) in milk. (D) MUFA/SFA ratio in milk. (E) Percentage of polyunsaturated fatty acids (PUFA) in milk. (E) Percentage of ω-3 fatty acids in milk. (F) Percentage of ω-6 fatty acids in milk. (G) ω-6:ω-3 ratio in milk. (n=10 dams).

Based on this finding, we assessed the specific fatty acid components of the milk fat using gas chromatography. These analyses showed a more desaturated and DHA-rich milk in the knockout (full results in Supplementary Table 3 and Supplementary Figure 3). At an aggregate level, knockout dams produced milk with 11% lower saturated fatty acids (SFA, Figure 4B, p=0.008), 12% higher monounsaturated fatty acids (MUFA, Figure 4C, p=0.009), but similar percentages of polyunsaturated fatty acids (PUFA, Figure 4E). The MUFA/SFA ratio showed that the knockout had a 24% higher level of desaturation (Figure 4D, p=0.004).

While PUFAs overall were similar between groups, knockout milk had 28% higher levels of ω-3 fatty acids (Figure 4F, p=0.013), driven primarily by a 42% increase in the ω-3 fatty acid docosahexaenoic acid (DHA; Supplementary Figure 3, p=0.031). There was a similar percentage of ω-6 fatty acids (Figure 4G), resulting in a 31% lower ω-6:ω-3 ratio (Figure 4H, p=0.008) in the milk of knockout dams. Individual ω-6 fatty acids did not significantly differ between groups (Supplementary Figure 3). Interestingly, the upstream precursors of DHA including alpha-linolenic acid (ALA) and eicosapentaenoic acid (EPA) were similar, suggesting that either the conversion from precursors into DHA may be increased or there is selective sparing of DHA from catabolism.

### Suppressed Expression of Adaptive Immune Markers and Increased Expression of Muscle Biosynthesis Genes

To understand the mechanisms by which adipocyte mTORC1 activation affects mammary gland biology, we performed bulk RNAseq on whole mammary gland explants from lactating wild-type and knockout dams. We identified 139 significantly differentially expressed genes between these groups (Figure 5A-B and Supplementary Table 1). In spite of the observed differences in milk fat and milk fatty acid composition, we were surprised that most fatty acid and triglyceride synthesis enzymes were similar between groups (Figure 5C). Several markers of adipogenesis and PPARγ were upregulated in the knockout mammary glands including *Plin4, Adipoq, Cav2*, and *Fabp4* (Figure 5D), consistent with the observed increase in mammary adipocyte numbers (Figure 2B). There were no detectable changes in PPARγ transcripts. We also identified several upregulated genes involved in ω-3 eicosanoid metabolism, including the enzymes *Cyp2e1, Gpx3, Ephx2*, and *Pla2g4a* (Figure 5E). Conversely, COX1 (*Ptgs1*), was significantly downregulated (Figure 5E).

**Figure 5:**
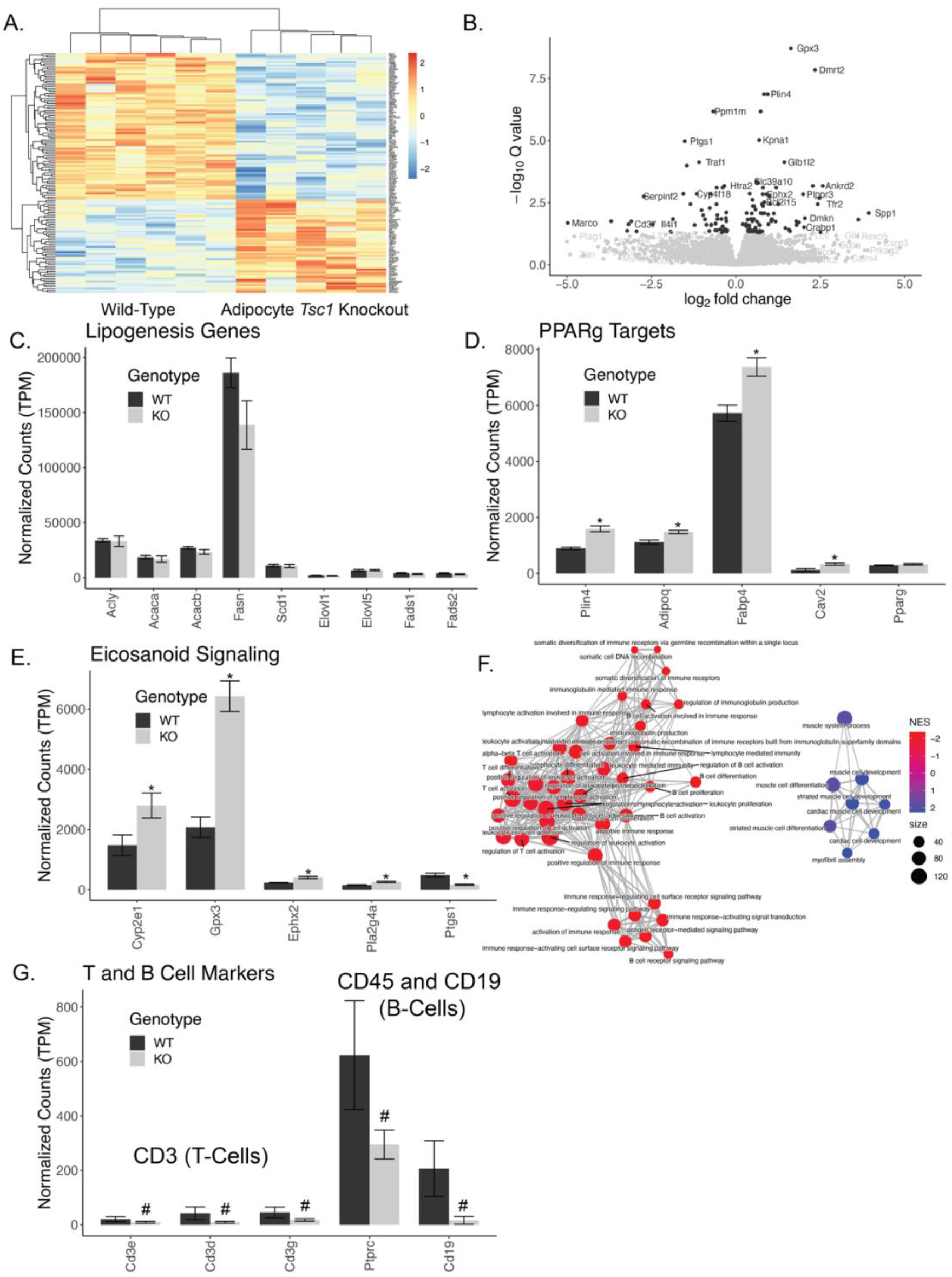
Transcriptomic analysis of mammary glands. (A) Heatmap of significantly differentially expressed genes between wild-type and knockout mammary glands. (B) Volcano plot comparing fold change to significance of specific genes. (C) Selected lipogenic gene expression. (D) Selected PPARγ target genes. (E) Selected genes involved in eicosanoid metabolism and signaling. (F) Network map of the top differentially expressed pathways from gene ontology – biological process. Jaccard distances were used to calculate similarity between pathways based on overlapping genes, and those distances are indicated as the edges. Net enrichment score (NES) indicate up or downregulation, while node size indicates the number of genes in that pathway. (G) T- and B-cell gene expression markers. Asterisks indicate q<0.05; number signs indicate p<0.05. (n=11 dams).

Despite the modest numbers of significantly differentially expressed genes, gene set enrichment analyses identified 220 significantly differentially expressed biological pathways (180 downregulated and 40 upregulated; Supplementary Table 2). By identifying genes in these pathways, they largely fell into two clusters of significantly differentially expressed pathways, one set related to the downregulation of adaptive immune differentiation and function, and another related to upregulation of striated muscle function (Figure 5F). To further explore the potential effects on adaptive immune cell function, we examined the expression of the T-cell marker genes encoding for CD3 and the B-cell marker genes encoding for CD45 and CD19.

Each of these markers were reduced 20-92% suggesting a potential reduction in adaptive immune cells in these mammary glands (Figure 5G). This could be primarily related to the increased DHA levels in the milk of knockout dams which could promote an anti-inflammatory state and reduce adaptive immune cell levels.

## Discussion

Milk fat is the most variable macronutrient in human milk and contributes the most to differences in energy content of milk (37). In this study, we show that hyperactivation of adipocyte mTORC1 via adipocyte-specific deletion of *Tsc1* alters milk fat composition and mammary gland adipocyte histology. Importantly, our approach is expected to activate mTORC1 in all adiponectin-expressing cells, including both peripheral and mammary adipocyte depots (24, 25). The positive role of mTORC1 in adipocyte biology has been well established. mTORC1 is necessary for adipocyte differentiation in peripheral adipose depots (38–40) but less is known about its role in mammary adipocyte differentiation and growth during lactation. It is interesting that we observed increased adipocyte numbers and elevated markers of adipocyte differentiation in adipocyte *Tsc1* knockout mammary glands, as the Adiponectin-Cre is not expected to be activated until late in differentiation. It is also worth noting that there is evidence of de- and re-differentiation of post-involution mammary adipocytes, but to avoid this concern, our study focused on mice in their first pregnancy. The increased adipocyte hyperplasia could suggest a signal promoting mammary adipogenesis derived from peripheral adipocytes.

mTORC1 is important for lipogenesis, and the chronic absence of mTORC1 activity in adipocytes (by *Raptor* ablation) results in lipodystrophic mice (41, 42). Gain of function studies (via tissue-specific *Tsc1* knockout) of mTORC1 in adipocytes resulted in increased *in vitro* palmitate esterification in inguinal adipose tissue (43). In our adipocyte *Tsc1* knockout model, we were surprised that there were no obvious increases in lipogenic enzymes in the mammary gland in spite of increased milk fat composition. This is consistent with data from *Raptor* knockout adipocytes, which have elevated (not decreased) ACC, ACLY, and FASN protein levels (41). We propose that in the knockouts there is increased peripheral synthesis of lipids, which are then transported to the mammary gland adipocytes for storage and secretion into the milk, though we cannot rule out non-transcriptional upregulation of lipogenesis in mammary glands. Our hypothesis of increased trafficking of lipids is consistent with elevated expression of the fatty acid transporter *Fabp4* (Figure 5C). Further studies using depot-specific activation of mammary adipocytes will be important to separate the roles of peripheral adipocytes from mammary adipocytes with respect to lactation.

Transgenic activation of Akt (an upstream activator of mTORC1) in mammary epithelial cells resulted in higher milk fat percentage during lactation and larger milk lipid droplet size compared to the control mice (44). In a separate study, supplementation of the mTORC1 activating branched chain amino acid, valine, increased mammary gland lipogenic activity during lactation in a way that was reversible by the mTORC1 inhibitor rapamycin (45). Together, these data suggest that mTORC1 activation may positively regulate milk lipids through multiple cell-types. We found that the secreted milk volume measured at PND10.5 was similar across genotypes. This suggests that the main driver of the modestly increased offspring weight could be the increase in milk fat percentage.

In addition to total lipids, we show an increase in both the relative desaturation of lipids, and the levels of DHA in milk of adipocyte *Tsc1* knockout mice. DHA is an essential ω-3 fatty acid important for infant growth and development and has been linked to improved cognitive performance, psychomotor development, and visual acuity (46–48). DHA and EPA levels are highly variable in human milk, and a better understanding of the physiological signals that control DHA levels in milk is important to optimize the delivery of essential lipids to the infant. We examined the expression of the phosphatidylcholine-DHA transporter *Mfsd2a* (49) but did not detect any differences in our mammary gland expression data. The increase in DHA levels may also be linked to our observation of reduced gene expression of markers of adaptive immune cells, as DHA has been shown to suppress the adaptive immune response (50). We show that several enzymes that convert DHA into bioactive lipids are upregulated in our lysates (Figure 5E). DHA-derived eicosanoids, such as D-series resolvins and protectins, could serve as negative signals to reduce the number of B and T cells in the mammary gland. This in turn could affect both mammary gland morphology and the secretion of antibodies into the milk.

This is the first report that adipocyte mTORC1 activation alters the lipids in milk of lactating mice, and provides important new data towards our understanding of lipid metabolism during a critical developmental window. There are several strengths to our approach, including the use of matched diets, single parity (to avoid mammary adipocyte de- and re-differentiation concerns), and normalized litter sizes to comprehensively evaluate milk lipids and mammary gene expression. However, there are several limitations to this approach including the inability to exclude *in utero* effects on offspring growth, the inability to separate the roles of peripheral and mammary adipocyte depots, and the lack of a clear mechanism by which mammary (or peripheral) adipocytes result in increased milk lipids, milk fat saturation, and milk DHA levels.

Our data show that elevations in mTORC1 activity in adipocytes of pregnant and lactating dams can impact milk composition, offspring weight, and mammary gland gene expression and morphology. These findings are crucial to better understand the effects of nutrient sensing in the mammary gland on milk production and offspring health. These data support our hypothesis that mTORC1 activation in adipocytes increases the mammary gland capacity to produce fat and secrete it into the milk. Future work will focus on the mechanisms by which mTORC1 could be influencing mammary gland function and milk secretion to address the effects of maternal excess nutrient sensing on lactation and infant health.

## Acknowledgements

This work was supported by funding from the NIH (R01DK107535 to DB and K01DK102526 to BG) and supported by core facilities at the Rogel Cancer Center (NIH P30CA046592), the MRC2 Metabolomics Core (NIH U24DK097153), and the Michigan Nutrition and Obesity Research Center (NIH P30DK089503).

## Author Contributions

This study was conceptualized by NEH, DB, and BG. Resources and funding were provided by BG and DB. The project was administered by DB and NEH. Investigation was performed by NEH, ACM, HH, JRR, ZC, and MCM. Formal analyses, computation, testing, and visualizations were performed by NEH, ACM, and DB. Data was curated by NEH. The initial draft was written by NEH and DB, with all authors reviewing and approving of the final version.

## Supplementary Figures

**Supplementary Figure 1:**
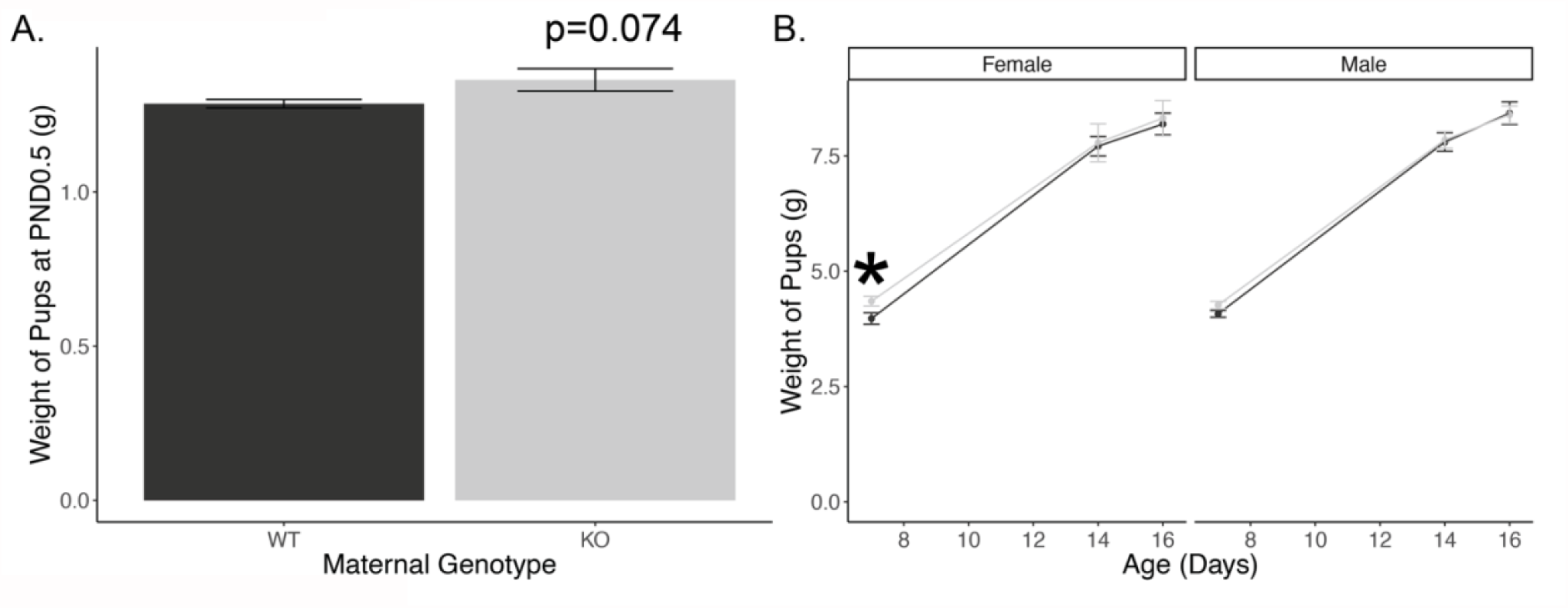
Offspring weight. (A) Weight of offspring at birth. (n=84 offspring from 11 dams). (B) Weight of offspring at PND7.5 through PND16.5. (n=43 offspring from 11 dams).

**Supplementary Figure 2:**
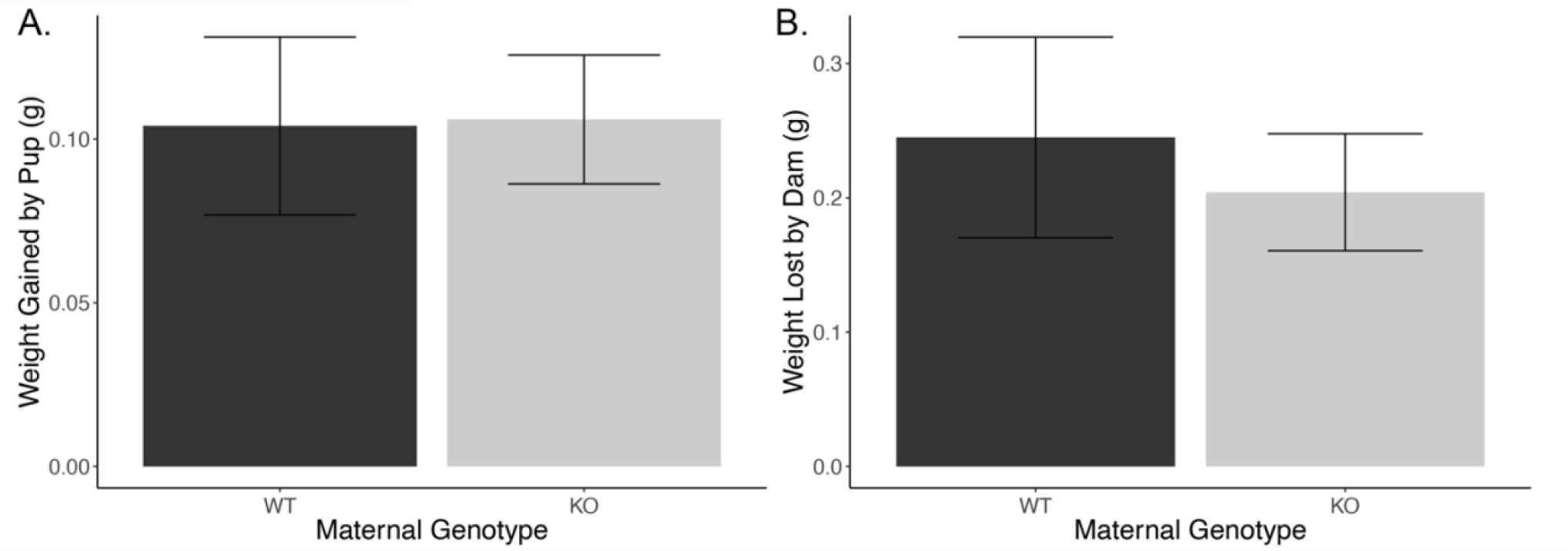
Milk volume measurement. (A) Weight gained by the offspring during 1h of nursing. (B) Weight lost by dam during 1h of nursing. (n=11 dams).

**Supplementary Figure 3:**
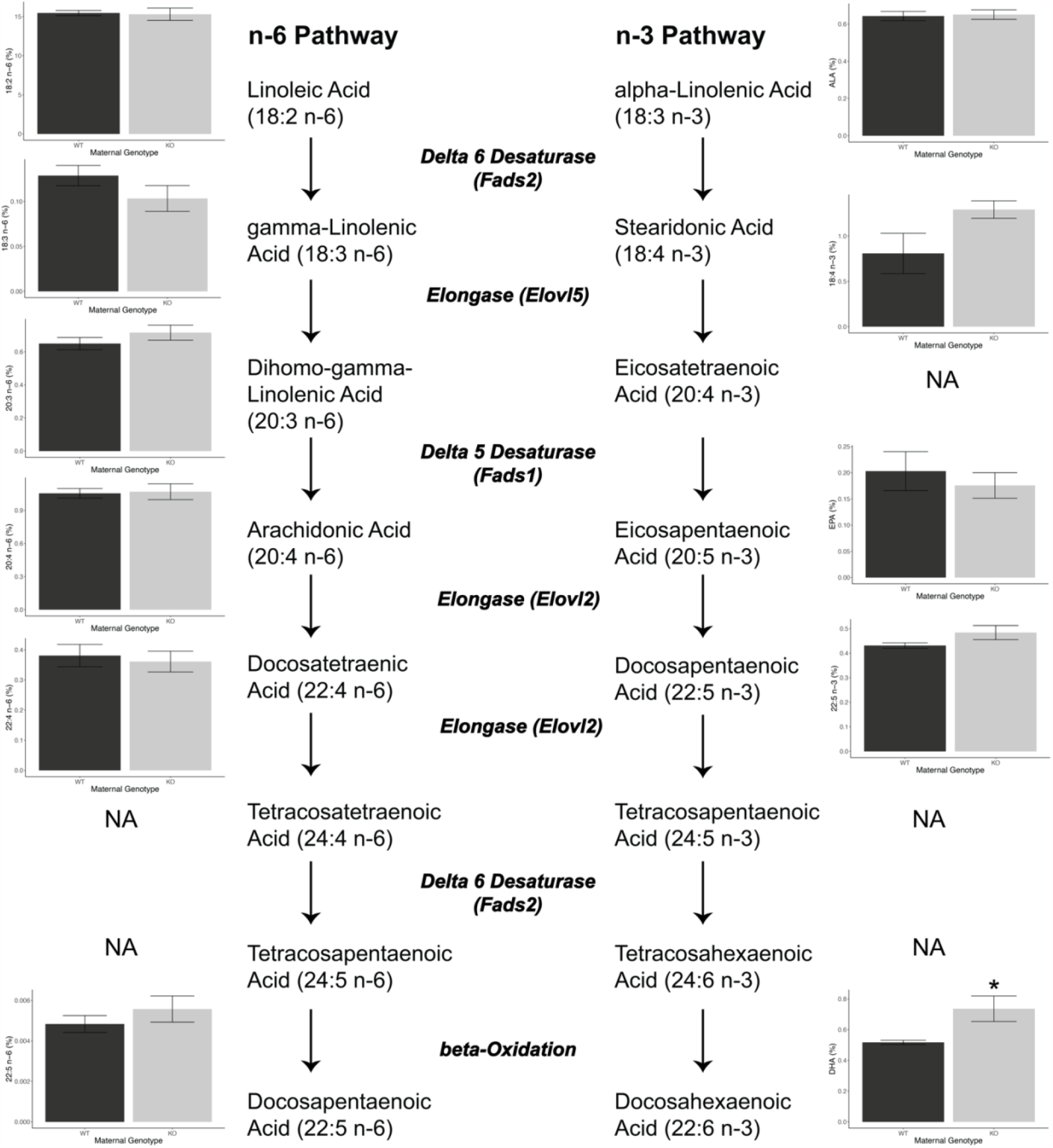
Fatty acid pathways. ω-3 and ω-6 metabolic pathways, enzymes, and detected ω-3 and ω-6 fatty acids. Fatty acids that were not detected are noted with NA. (n=10 dams).

***Supplementary Table 1: Complete gene expression data***. Gene level expression data including nominal (pvalue) and FDR-adjusted (padj) p-values. NA indicates that the gene was not evaluated by DESeq2, generally due to low expression.

***Supplementary Table 2: Gene set enrichment analyses using gene ontology – biological pathways***. Gene-ontology pathways that reached statistical significance. NES indicates the net enrichment score with positive indicating upregulation of the pathway. Core enrichment indicates the specific genes that drive significance of this pathway.

***Supplementary Table 3: Milk fatty acid composition***. Group mean and standard errors are presented along with the percent change relative to milk of wild-type. Test indicates the pairwise test (based on tests of normality and heteroscedasticity) and the resultant p-value.

